# KAT5 Acetylates Human Chromatin Protein PC4 to promote DNA Repair

**DOI:** 10.1101/2024.01.12.575390

**Authors:** Sweta Sikder, Aayushi Agrawal, Siddharth Singh, Ramalingam Peraman, Viswanathan Ravichandran, Tapas K. Kundu

**Author notes:** **Address for correspondence**: Tapas K Kundu,. Authors contributed equally to this manuscript (Sweta Sikder, Aayushi Agrawal).

## Abstract

Human positive coactivator 4 (PC4), is a highly abundant non-histone chromatin protein involved in diverse cellular processes, including transcription regulation, genome organization, DNA repair, etc. The majority of PC4 exists in a phosphorylated state in cells, which impinges its acetylation by p300 and thereby inhibits its double-stranded DNA binding ability and transcriptional co-activator function. Recently, we have shown that PC4 interacts with linker histone H1 in its phosphorylated state, and this interaction is important for PC4- mediated chromatin compaction. PC4 was also found to be an activator of non-homologous end joining and DSB repair activity. Knockdown of PC4 causes drastic decompaction and enhanced autophagy in the cells. Mechanistically, in the absence of PC4, the genome becomes highly vulnerable to DNA damage with an altered epigenetic landscape. Here, we report that other than p300, PC4 also gets acetylated by DNA repair facilitating lysine acetyltransferase KAT5 (Tip60), at K80 residue when the cells are subjected to DNA damage. The vulnerability of DNA in PC4 devoid cells could be substantially reduced by reintroducing wild-type PC4 to the cells but not the mutant PC4 (K80R PC4), defective in KAT5-mediated acetylation. Presumably, KAT5-mediated acetylation of PC4 at K80 residue facilitates the DNA repair machinery at the damage site and thus contributes to DNA damage repair, a process that could be of high significance both in cancer and in aging.

## 1. Introduction

The human genome is under constant threat from various genotoxic agents that can cause severe damage to our genetic material, DNA. Erroneous or inefficient repairing of the genome upon such genotoxic insults can cause changes in the DNA sequence itself, which if not corrected can trigger genomic instability and thereby lead to a harmful cellular environment and ultimately cell death. However, as a safeguard mechanism, cells counteract these adverse effects by activating the DNA damage response (DDR), which causes a synchronized series of events that regulates cell cycle progression and repair of DNA lesions. The tight packaging of the genomic DNA very often poses a threat to the DNA repair pathway in living cells. Therefore, cells modulate the chromatin structure at the site of damage with the help of various factors such as post-translational histone modifications, ATP-dependent chromatin remodelers, etc. These factors in concert help the DNA repair machinery to access the DNA lesions.

In addition to the above-mentioned factors, various non-histone chromatin associated proteins also play an important role either by directly interacting with the DNA repair machinery proteins or by regulating the chromatin structure [1]. Histone PTMs constitute an important layer of epigenomic information and some of these modifications have been shown to regulate DSB repair responses. One such modification is H3K9me3 which is enriched in constitutive heterochromatin regions and is shown to increase at damaged site, both in heterochromatin and euchromatin regions [2–4]. The players involved in H3K9me3 pathway, such as writers (SUV39H1/2, SETDB1) and readers (HP1, TIP60) were shown to promote DSB repair by Homologous Recombination (HR) pathway.

In eukaryotic chromatin, about 10% of all nucleosomes consist of histone H2A variant H2AX instead of canonical H2A. Previous studies have shown that, following the induction of DSBs, the MRN complex (MRE11–RAD50–NBS1) binds to broken DNA ends and recruits ATM, ATR, and/or DNA protein kinase (the PIK-family protein kinases), resulting in the initial phosphorylation of H2AX [5, 6]. The adaptor protein MDC1 then associates with phosphorylated H2AX (γ-H2AX) and recruits additional activated ATMs to the sites of DSBs [7]. This positive feedback loop leads to the expansion of the γ-H2AX region surrounding DSBs, to up to 2 Mb, and provides docking sites for other DNA damage and repair proteins, including 53BP1 and BRCA1.

Tip60 plays a crucial role in DSB repair and its inactivation leads to increased sensitivity to ionizing radiation and increased levels of chromosomal aberrations [8–12]. Tip60 is also required for the activation of the ATM kinase which phosphorylates multiple DNA damage response proteins, including nbs1, p53, chk2 and SMC1 in response to DSB detection [13]. Further, Tip60 is a haplo-insufficient tumor suppressor, and its aberrant expression has been detected in various cancers [14–16]. These studies demonstrate that a key function of Tip60 is to protect cells from genomic instability and to suppress potentially transforming events which can lead to cancer.

Human positive coactivator 4, PC4 has been reported to get recruited early at DNA damage site [17]. It has been shown to interact with the DNA repair protein XPG and gets recruited to DNA damage site (bubbled DNA). PC4 also prevents mutagenesis and killing by oxidative DNA damage repair by its ssDNA binding ability [18]. Delving more into the molecular repair pathway, it has been shown to enhance joining of non-homologous end joining repair and double strand break repair activity and also in homologous recombination repair [19, 20]. Interestingly, PC4 interacts with the heterochromatin HP1α, in vivo [21]. In this study, we pose the question that whether the DNA repair activity of PC4 is being modulated by any post translational modifications. Interestingly, we found that the DNA damage responsive acetyltransferase, KAT5/Tip60 acetylates PC4 at specific sites, in the cellular context. The Tip60-mediated acetylation of PC4 was found to be critical for DNA damage repair, which is presumably linked to breast cancer pathobiology [22].

## 2. Materials and Methods

### 2.1. Filter Binding Assay

1µg of Histone H4 or Histone H3 was incubated with equivalent activity of baculovirus expressed full length human Tip60 and p300 at 30°C for 30 minutes in 2X KAT buffer (1X composition: 50mM Tris-HCl pH 8.0, 10% glycerol, 1mM DTT, 1mM PMSF, 0.1mM EDTA), 10mM Na-butyrate and 1µl of 2.1 Ci/mmol of [3H]-acetyl CoA. The reaction mixture was then spotted on a Phosphocellulose P-81 filter paper. The radioactive counts were recorded on a Wallac 1409 Liquid scintillation counter.

### 2.2. Gel Assay

KAT assays were performed using 500ng/1μg of Histone H3 or 500ng/1μg of recombinant PC4, or 500ng/1μg of p53 incubated in HAT assay buffer at 30°C for 30 minutes with or without baculovirus expressed recombinant Tip60, 1µl of 3.3 Ci/mmol of 3H-acetyl CoA and 10mM Na-butyrate. To visualize the radiolabeled acetylated protein, the reaction products were resolved electrophoretically on 12% SDS-polyacrylamide gel and subjected to fluorography. The gel was stained by coomassie to ascertain the presence of protein in equal amounts in each of the reaction and was later dehydrated in DMSO for 1 hour. Later the gel was incubated in scintillation fluid (PPO solution in DMSO) for 30 minutes and then rehydrated in water for 2hrs. The gel was dried using a gel drier and exposed in an X-ray cassette using a film for 7days in -80⁰C cooler. The film was later developed to get intensity profiles for each of the reaction.

### 2.3. Acetylation of PC4

Mass acetylation of PC4 was carried out using 1µg of His-PC4. Reaction mixture containing recombinant Tip60 (∼ 30,000 Counts/µl), 2X KAT buffer, 10mM Na-Butyrate and 1.68mM acetyl-CoA was incubated at 30°C for 30 minutes, followed by replenishment with the enzyme and Acetyl CoA every 1 hour and then kept for 6-8 hours after the final replenishment. Mock acetylation reaction was also set up without the enzyme.

### 2.4. Mass spectrometric analysis

#### 2.4.1. Preparation of the sample

1µg of PC4 was mass acetylated as given in the previous section. The acetylated sample was electrophoresed in a 12% SDS PAGE and was silver stained by fixing in 40% methanol and 10% glacial acetic acid in water overnight. The gel was then rinsed thrice in 50% ethanol in water. The proteins were sensitized with 0.02% sodium thiosulfate for 2 minutes then rinsed in deionized water for a minute. The gel was then incubated with 0.1% silver nitrate for 20 minutes at room temperature. The gels were then rinsed thrice with deionized water. The gel was developed in freshly prepared 0.04% formalin/2% sodium carbonate (∼4-20 minutes). The reaction was stopped in 1% glacial acetic acid. The gel was imaged and then stored in 1% glacial acetic acid. The specific band for PC4 was cut and dried and given for mass spectrometric analysis.

#### 2.4.2. Mass spectrometric analysis of *in vitro* acetylated PC4

The mass spectrometric service of ITSI Biosciences, USA was availed to carry out the PTM analysis. In-gel digestion of the PC4 band was performed using trypsin as the enzyme. After trypsinization, peptides were transferred to another tube and two more extraction of peptides was done using 1:2 (v/v) 5 % Formic acid and acetonitrile. Eluates were dried down using a speed vac and reconstituted in 2% Acetonitrile with 0.1% Formic acid. Samples were then loaded onto a PicoFrit C18 nanospray column (New Objective) using a Thermo Scientific Surveyor Autosampler operated in the no waste injection mode. Peptides were eluted from the column using a linear acetonitrile gradient from 2 to 30% acetonitrile over 90 minutes followed by high and low organic washes for another 20 minutes into an LTQ XL mass spectrometer (Thermo Scientific) via a nanospray source with the spray voltage set to 1.8kV and the ion transfer capillary set at 180°C. A data-dependent Top 5 method was used where a full MS scan from m/z 400-1500 was followed by MS/MS scans on the five most abundant ions and multistage activation was turned on for detecting acetylated peptides. Raw data files were searched using Proteome Discoverer 1.3 (Thermo Scientific) and the SEQUEST algorithm against the most recent species-specific database for Homo sapiens and custom database with the protein of interest from UniProt. Trypsin was the selected enzyme allowing for up to three missed cleavages per peptide; Carbamidomethyl Cysteine was used as a static modification, Acetylation on lysine and Oxidation of Methionine as a variable modification. Proteins were identified when two or more unique peptides had X-correlation scores greater than 1.5, 2.0, and 2.5 for respective charge states of +1, +2, and +3 (Fig. S1).

#### 2.4.3. Pull down assay

Cells were cultured in dishes. After 24 hours, cells were washed with 10 ml of pre-chilled PBS and lysed for 60 minutes in 1 ml RIPA lysis buffer (150mM NaCl, 60mM Tris–HCl, 0.5 mM EDTA, 10% NP-40, pH 7.4) with 1X of protease inhibitor cocktail (Sigma). Next, 1 µg of antibody or IgG was used to bind to the bait proteins for overnight binding at 4° on rotor, and then incubated with 30 µL protein G for an additional 5-6 hours. Finally, the protein G was washed with 1 ml of RIPA lysis buffer three times at 4°C at 1500 rpm for 2 minutes and then heated for 10 minutes at 95◦C with 30 µL of 2× SDS loading buffer. Samples were analysed by SDS-PAGE and western blotting.

#### 2.4.4. Mass spectrometric analysis of PC4 and acetylated PC4 interacting proteins

The pull-down complexes were then resolved on a 12% SDS PAGE. Each lane of the SDS-PAGE gel was excised into five equally sized segments. Gel pieces were processed using the following protocol: (1) Destained with 50% methanol containing 0.1% acetic acid and washed with 50 mM ammonium bicarbonate followed by acetonitrile; (2) Reduced with 10 mM dithiothreitol at 37°C for an hour followed by alkylation with 55 mM iodoacetamide at RT for 45 minutes; (3) Digested with trypsin (Thermoscientific) at 37 ◦C for 4 h; (4) Quenched with formic acid and the supernatant was analysed directly without further processing. Each gel digest was analysed by nano LC/MS/MS with a RSLC Nano system (Thermoscientific Ultimate Dionex 3000), MS system (Thermoscientific Orbitrap Exploris 240). Peptides were loaded on a trapping column and eluted over a 75µmX25cm analytical column. Data interpretation was done using Proteome Discoverer, Version 2.5.

### 2.5. Generation of Polyclonal Antibody (PC4 K80Ac)

Polyclonal antibodies were generated by injecting the Keyhole Limpet Hemocyanin (KLH) coupled peptides in rabbits. New Zealand strains of rabbits were primed with an emulsion of the peptide-KLH conjugate in complete adjuvant. Before the priming, a few ml of blood was collected from the rabbit (control serum). Booster doses were given every 2-3 weeks till the serum showed sufficient strength and specificity against the target protein or modification. Major bleed of 15-20 ml was collected twice in the course of raising the antibody. Serum was separated from the collected blood and IgG was purified using Protein G Sepharose beads. The IgG was eluted from the beads using 100 mM glycine, pH 3.0 and collected into tubes containing Tris-Cl buffer to neutralize the acidic pH of the elution buffer. The IgG purified was then dialyzed against 50% glycerol in PBS. A peptide was generated spanning K80Ac site in PC4 and was used to raise K80Ac specific polyclonal Ab of PC4. The peptide was KLH conjugated KGK (ac) VLIDIREYWMDC (Fig. S4).

### 2.6. Comet assay

The comet assay, or single cell gel electrophoresis assay (SCGE), is a common technique for measurement of DNA damage in individual cells. Under an electrophoretic field, damaged cellular DNA (containing fragments and strand breaks) is separated from intact DNA, yielding a classic “comet tail” shape under the microscope. Extent of DNA damage is usually visually estimated by comet tail measurement. Briefly, 0.8% low melting point (LMP) agarose was prepared in saline and was maintained at 39 ◦C to prevent solidification. Subsequently, individual cells (around 20,000 cells after 48 hours of transfection with Flag constructs of PC4 and mutant PC4) were mixed with molten agarose in 1:10 (v/v) ratio. The resulting suspension was layered onto the frosted slides. The slides were placed on ice for approximately 5 min to allow the agarose to solidify. Subsequently, slides were immersed in lysis solution (2.5 M NaCl, 100 mM EDTA with fresh 1% Triton-X-100 and 10% DMSO) for 1 hour to eliminate non-nuclear components. The slides were further immersed in alkaline buffer (300 mM NaOH, 1 mM EDTA, pH = 13) for 20 min to allow the DNA to unwind and to convert alkali labile sites to single strand breaks. Electrophoresis was conducted for 30 min at 15V and 200 mA (at a rate of 0.6V/cm) using a compact power supply. The slides were gently washed with 0.4 M Tris (pH = 7.5) to remove alkali and detergents. The cells were stained with propidium iodide (PI – 20 g/ml) and were visualized by epifluorescence microscopy. Under these conditions, the damaged DNA (containing cleavage and strand breaks) migrates further than intact DNA and produces a “comet tail” shape. Approximately 100-150 images per slide were captured from different imaging fields and were analyzed with the appropriate software.

### 2.7. Immunofluorescence microscopy

Cells grown on poly-L-lysine-coated cover slips were permeabilized by 1% Triton X-100 in PBS for 10 minutes, followed by blocking in 1% fetal bovine serum for 45 minutes. Probing was done with anti-PC4, anti-K80ac PC4, anti-γ.H2AX (CST), and anti-Tip60 polyclonal antibodies for 1 hour at room temperature followed by secondary antibodies conjugated to Alexa488 and Alexa 568 (Invitrogen). The cells were stained with 0.1 μg/ml of DAPI (Sigma) in PBS for 5 minutes to visualize the nuclei. Mounting was done with 70% glycerol Fluorescence images was taken through confocal canning microscope LSM510 META and ZEN LSM810.

### 2.8. Cell culture and transfection

HEK 293 and ZR-75-1 cells were obtained from ATCC. HEK293 and ZR-75-1 cells were cultured in DMEM and RPMI 1640 medium respectively, supplemented with 10% fetal bovine serum at 37◦C with 5% CO2. HEK293 cells were transfected using Lipofactamine 2000 according to the manufacturer’s instructions (Invitrogen).

### 2.9. Stable cell line generation

HEK 293 cells constitutively expressing shRNA against the ORF of Tip60 (Clone ID: V2LHS_282664, Dharmacon Pgipz shRNA; Mature sequence: TTCCATCAGAGCTGTCCTG) mRNA was established to make a PC4 and Tip60 knockdown cell line respectively. To neglect the effect of shRNA transduction a cell line expressing a non-silencing shRNA control was also established. The non-silencing control hairpin sequence is as follows: 22mer sense: ATCTCGCTTGGGCGAGAGTAAG 22mer antisense: CTTACTCTCGCCCAAGCGAGAG This sequence does not match any known mammalian genes (had at least 3 or more mismatches against any gene as determined via nucleotide alignment/BLAST of 22mer sense sequence). These cell lines were generated using 10 μg pGIPZ lentiviral shRNAs targeting PC4 and helper plasmids (5 μg psPAX2, 1.5 μg pRSV-Rev, 3.5 μg pCMV-VSV-G). 10 μg of sh-plasmid was mixed with helper plasmids (5 μg psPAX2, 1.5 μg pRSV-Revs, 3.5 μg pCMV-VSV-G) and were co-transfected into HEK293T cells using the calcium phosphate method. 48 hours post-transfection media containing assembled virus was collected and its titre was estimated. Desired cell line (here HEK 293) was infected with 105 IU/ml virus. Infected cells were subjected to selection pressure 72 hours post-transfection. Cells were first sorted for positive GFP signals and the GFP sorted cells were grown to establish the stable cell line. To validate the extent of knockdown, Tip60 levels were checked at the protein level and compared against the non-silencing shRNA harbouring HEK 293 cells.

### 2.10. Statistical analysis

The GraphPad Prism software (California, USA) was used to conduct all statistical analysis. All statistical results are presented as the mean ± SD or mean ± SEM, as indicated. P values (* P < 0.05, ** P < 0.01, *** P < 0.001, **** P < 0.0001) were obtained using Student’s t test (two-tailed) or one/two-way ANOVA from multiple independent biological replicates.

## 3. Results

### 3.1. PC4 is a substrate of KAT5 (Tip60) acetyltransferase

Owing to its role in DNA damage, we wanted to investigate PC4 as a potential substrate of the DNA damage responsive acetyltransferase, KAT5 (Tip60). The bacterially expressed untagged recombinant human PC4 was subjected to *in vitro* acetylation by both p300 and Tip60 (baculovirus expressed full-length recombinant His-tagged). p53, a known substrate of Tip60 [23], was taken as a non-histone protein positive control apart from the recombinant histone H3. The reaction mixtures containing equivalent activities of either p300 or Tip60 (as normalized by filter binding assay using Histone H3 and H4 respectively) and the indicated amount of proteins (Fig. 1A) were incubated with [^3^H] acetyl-CoA for 30 minutes at 30°C. The reaction mixtures were analyzed by autoradiography upon resolving on SDS-PAGE gel. It was observed that PC4 was not only acetylated by p300 but also with equivalent activity of Tip60 (Fig. 1A, middle panel). Thus, p300 is not an exclusive acetyltransferase which can acetylate PC4 *in vitro*. We thus show that, like p53 (Fig. 1A, right panel), PC4 might be also a substrate of the acetyltransferase Tip60. Once PC4 was established as a substrate of acetyltransferase of KAT5 (Tip60) by *in vitro* acetyltransferase assay, we next wanted to identify the specific acetylation sites through mass spectrometric analysis (as described in materials and methods). PC4 was mass acetylated and then subjected to mass spectrometric analysis for identification of the residues in PC4 which could be acetylated. Our mass spectrometric PTM analysis revealed confident acetylation sites in PC4 (Fig. 1B and S1). We narrowed down to 4 high scoring acetylated residues (represented in green) in the protein PC4 as indicated in Fig. 1B. The table shows the change in the mass of the peptides due to an acetylated lysine residue obtained by mass spectrometric analysis.

**Figure 1.**
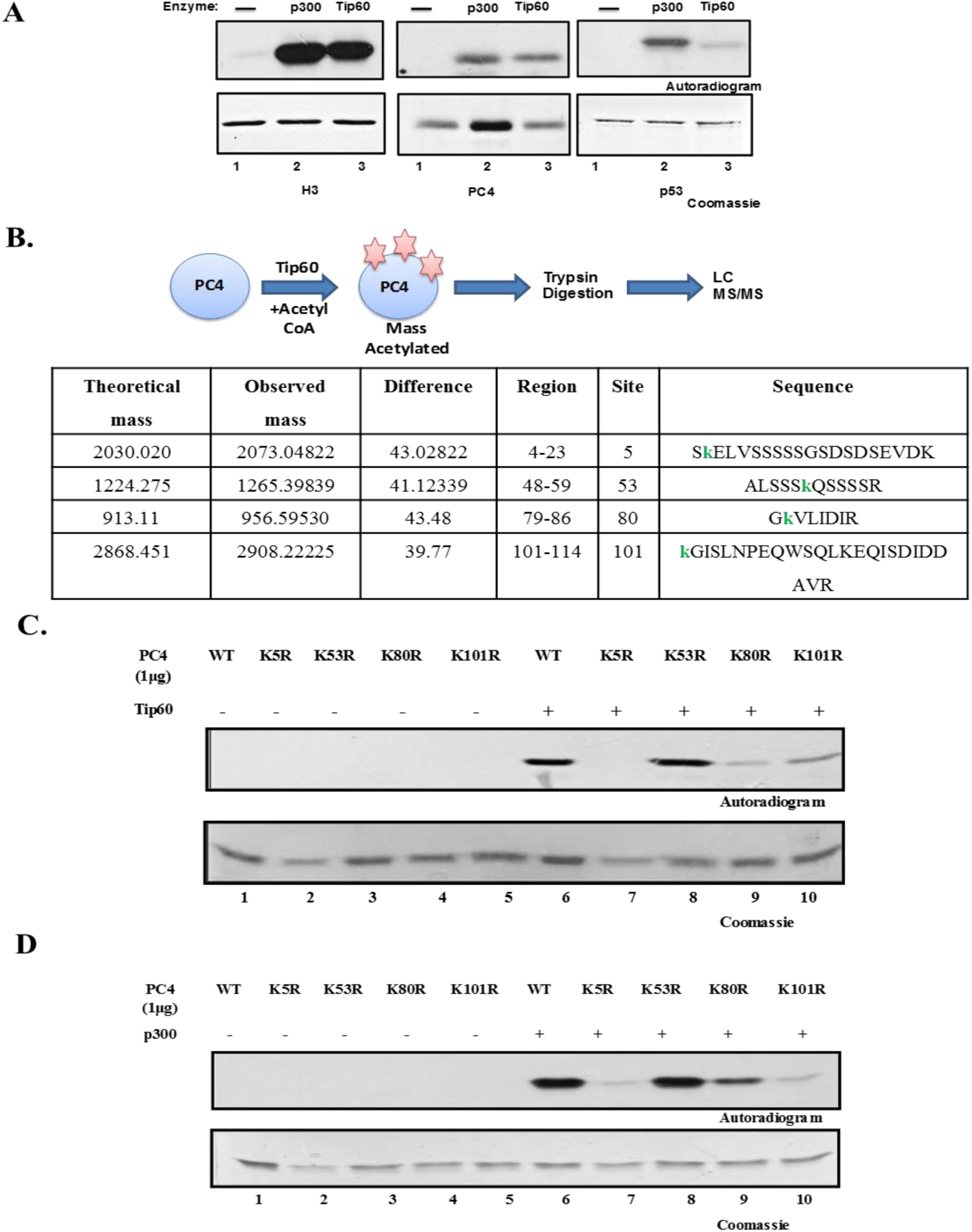
KAT5 acetylates PC4 *in vitro* at K80 residue specifically. (A) Purified proteins were incubated with [^3^H] acetyl-CoA for 30 minutes at 30°C and then separated by SDS- PAGE (12%) and visualized by fluorography. 500ng of purified Histone H3(left panel) without any KAT (lane 1), with p300 (lane 2), and with Tip60 (lane 3) and bacterially expressed His-PC4 (middle panel) without any KAT (lane 1), with p300 (lane 2), with Tip60 (lane 3) p53 (right panel) without any KAT (lane 1) with p300 (lane 2), and with Tip60 (lane 3) were incubated. (B) *In vitro* mass acetylated PC4 was subjected to mass spectrometric analysis to determine the potential acetylated residues. Table shows representative data from mass spectrometric analysis showing the change in mass of peptides containing a single acetylated lysine residue. The lysine residue which was found to be acetylated is represented in small font in green. (C) KAT assay was performed with 1μg PC4 wild type (WT) and 1μg of acetylation site mutants K5R, K53R, K80R and K101R in the presence (lanes 6-10) or absence of Tip60 (lanes 1-5). (D) KAT assay was performed with of PC4 wild type (WT) and with acetylation site mutants K5R, K53R, K80R, K101R in presence (lanes 6-10) or absence of p300 (lanes 1-5).

### 3.2. Acetylation of PC4 by Tip60 is abrogated by mutation at K80 residue

Taking cue from the mass spectrometric data, we wanted to further validate the acetylation sites in PC4 by generating acetylation defective mutants. Single site mutants were created for each of the lysine sites (changed to arginine) shown by site directed mutagenesis (Fig. S2). *In vitro* acetylation assay with these acetylation defective mutants was carried out. Acetylation of PC4 K5R, K80R and K101R were highly compromised as compared to PC4 Wild type (WT) (Fig. 1C, lane 6 vs 7, 9 and 10 respectively). PC4 K53R showed no change in acetylation with Tip60 as compared to the wild type PC4 (Fig. 1C, lane 8). This data suggests that the K53 residue might not be a major site of acetylation in PC4 mediated by Tip60. Mock acetylation controls were used (Fig. 1C, lanes 1-5) where except the enzyme all components including protein were taken. These data further establish the fact that the acetylation observed is specifically mediated by the enzyme Tip60.

*In vitro* KAT assay with PC4 acetylation defective mutants was carried out in a concentration dependent manner to negate the effects of protein saturation and also to further confirm the mutants which were defective to acetylation by Tip60. Single mutation of lysine to arginine at 5th amino acid position of PC4 (PC4K5R) completely abolished its ability to get acetylated. Even at higher concentration no band was observed. (Fig. S3A, lane 6 and 7). PC4K80R and PC4K101R also showed compromised acetylation ability which was not affected by the different concentrations of the protein used (Fig. S3B, lanes 6-9). Mutation at K53 to arginine did not affect the acetylation ability of PC4 by Tip60 (Fig. S3A, lanes 8 and 9). Thus, lysine residues at 5, 80, and 101 appeared to be important for acetylation of PC4 by KAT5.

PC4 is also a substrate of acetyltransferase p300 (as shown in Fig. 1A). To further validate the lysine residues in PC4 identified as substrates of acetyltransferase Tip60 and not p300, KAT assay was performed with each acetylation site mutants and p300. Comparing Fig. 1C and 1D, PC4 K5R showed reduction in acetylation by both p300 (Fig. 1C, lane 7) as well as Tip60 (Fig. 1D, lane 7) whereas PC4 K53R showed no drastic change in acetylation by either p300 (Fig. 1D, lane 8) or Tip60 (Fig. 1C, lane 8). PC4 K101R also showed compromised acetylation ability both by p300 (Fig. 1D, lane 10) and Tip60 (Fig. 1C, lane 10). However, PC4 K80R shows a drastic reduction in Tip60 mediated acetylation (Fig. 1C, lane 9) but when subjected to acetylation by p300 (Fig. 1D, lane 9), it showed considerable amount of acetylation. Thus, PC4 K80 might be a putative site of acetylation in PC4 specific for Tip60. The highly conserved K80 residue of PC4 dwells in its C-terminal domain which is essential for its single strand DNA binding activity which in turn is critical for its DNA repair function [19, 20].

### 3.3. PC4 gets acetylated at K80 upon DNA damage conditions

To understand the physiological role of KAT5/Tip60 mediated acetylation of PC4, we raised a site specific (K80Ac) polyclonal antibody to detect the in vivo acetylated PC4. The polyclonal antibody was tested for its specificity and sensitivity for Tip60 acetylated PC4 at K80 site (Fig. S4). Considering the role of Tip60 in DNA damage and repair that has well been characterized [24, 25], we looked for PC4 acetylation at K80 sites under conditions of Cisplatin and Actinomycin D treatment. Actinomycin D is a DNA damage inducing agent which mostly forms DNA intercalation producing radical oxygen species and thereby causing genotoxic insult to the DNA. It has also been shown to induce the accumulation of γ-H2AX which forms complex with DNA repair proteins in the cells [26]. Cisplatin interacts with DNA and stalls cell proliferation, thereby activating DNA damage response. Thus, both the small molecules were administered to mimic a DNA damage state in the cells. γH2AX level was checked by western blot analysis to validate the *in vivo* conditions. We find elevated level of γ-H2AX which is known to be upregulated upon DNA damage as compared to H2A levels (used as an internal loading control). The cellular PC4 K80Ac mark was found to be enhanced upon treatment with Cisplatin and also with Actinomycin D as revealed by immunofluorescence and western blot analysis with the PC4K80Ac antibodies (Fig. 2). Interestingly, we observed that upon DNA damage, in the PC4 knock down cells, the residual PC4 also gets acetylated at the K80 site (Fig. 2E, panel 3, lanes 2,4,6). Collectively, our data suggest that PC4 acetylation at K80 site is responsive to DNA damage condition. Considering the role of PC4 in DNA damage repair, the acetylation by the Tip60 might play a crucial role in damage repair process.

**Figure 2.**
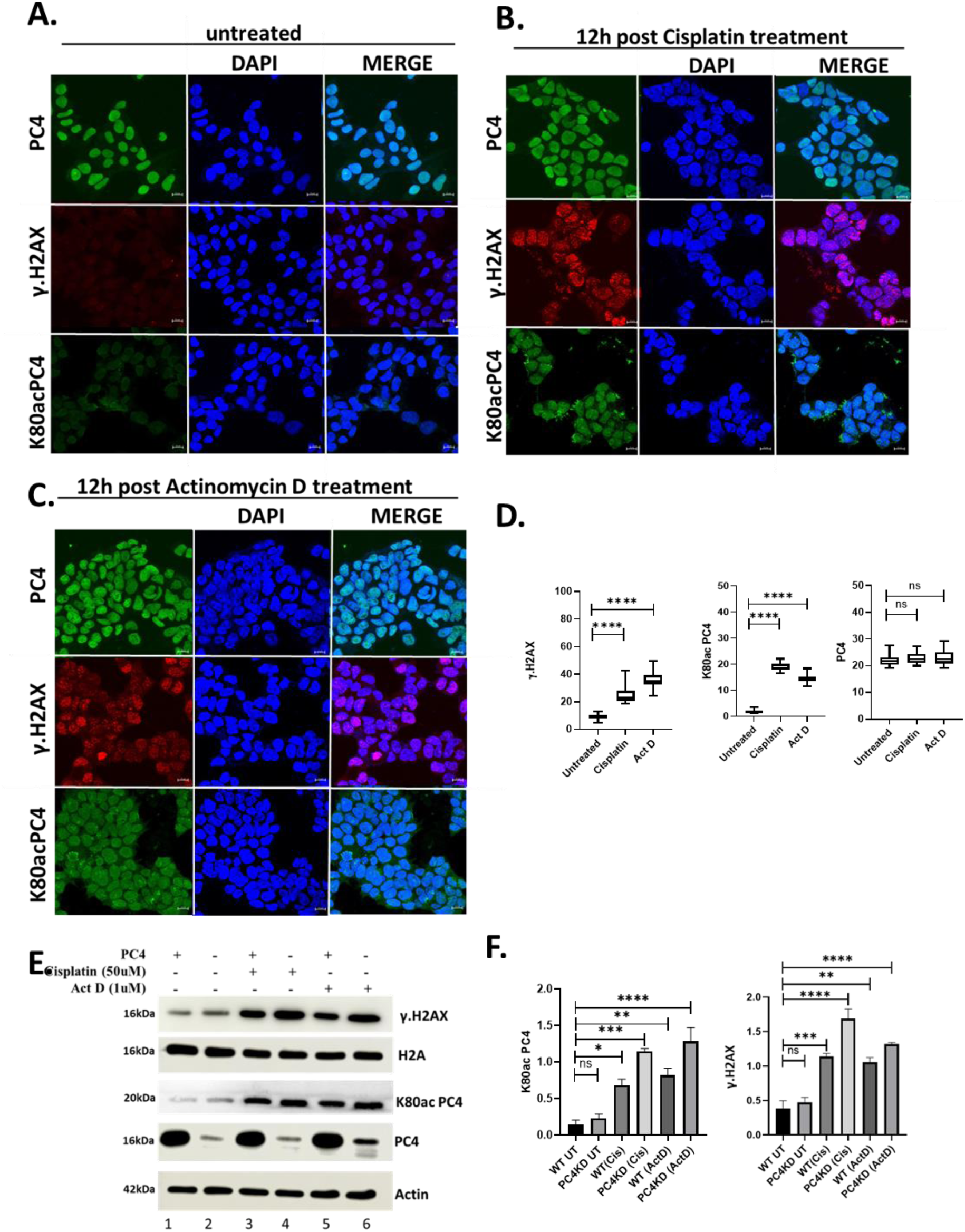
PC4 gets acetylated at K80 under DNA damage conditions. HEK293 cells were treated 0.2µM Actinomycin D and 30 µM Cisplatin for 12hours. Cells were stained after the aforementioned time point for treatment. Staining with K80Ac PC4 antibody was used to detect the acetylation levels, while PC4 antibody was used to detect the unmodified form of PC4. γ.H2AX was used as a DNA damage marker. Immunofluorescence images showing PC4, K80Ac PC4 and γ.H2AX levels in untreated (A), Actinomycin D treated (B), and Cisplatin treated (C) cells. (D) Fluorescence intensity was quantified using ImageJ and plotted (n=50 cells). (E) Immunoblotting with RIPA lysates were done using the specific antibodies. Equal Volumes of RIPA lysates were electrophoresed on a 12%SDS gel and probed with specific antibodies. The levels of H2A and actin was checked as a loading control for γ.H2AX and K80ac PC4 respectively. The figure is representative of three biological replicates and the quantified intensity values are plotted as shown in (F). Bars represent mean ± SE. Statistical analysis was carried out by one-way ANOVA using Dunnett’s test for multiple comparisons, \*\*\*\**p* < 0.0001, \*\*\**p* < 0.001, \*\**p* < 0.01 and \**p* < 0.01. (Scale Bar- 10µm).

After establishing the cellular condition at which Tip60-mediated acetylation of PC4 at K80 site gets enhanced, we wanted to investigate the nuclear localization of this specific acetylation. Mammalian Flag-tagged expression constructs of both the wild type and the acetylation defective mutant (K80R) were created. To check the expression and also their cellular localization, immunofluorescence studies were carried out. Flag constructs of both wild-type and mutant PC4 were transfected in PC4 knockdown cells. Mutation at K80 site to arginine did not perturb its cellular localization as both wild type and mutant protein showed similar nuclear localization (Fig. S5).

### 3.4. Tip60 is the bonafide acetyltransferase to acetylate PC4 at K80

Although in vitro acetylation assays showed that K80 site of PC4 is indeed acetylated by Tip60, these data do not ensure that in the cellular context or physiologically, Tip60 is involved in the acetylation of PC4 at K80. In order to establish the role of Tip60 in the acetylation of PC4 in cells, we generated the stable knockdown of Tip60 in HEK 293 cell line (sh-Tip60) using a lentiviral mediated short hairpin RNA (shRNA) delivery system. The sh- Tip60 cells showed Tip60 downregulation both at transcript and protein level (Fig. S6 and 3A respectively).

We observed that when cells were treated with Actinomycin D, PC4 got profusely acetylated at K80 site, however, in absence of Tip60 (in Tip60 knockdown cells) there was a drastic decrease in the acetylation level, as expected (Figure 3B, top panel). Interestingly, it was found that upon DNA damage by Actinomycin D, the γ-H2AX level, is substantially increased in the Tip60 knockdown cells, suggesting the accumulation of damage, but not the acetylated PC4, in absence of Tip60 (Fig. 3B).

**Figure 3.**
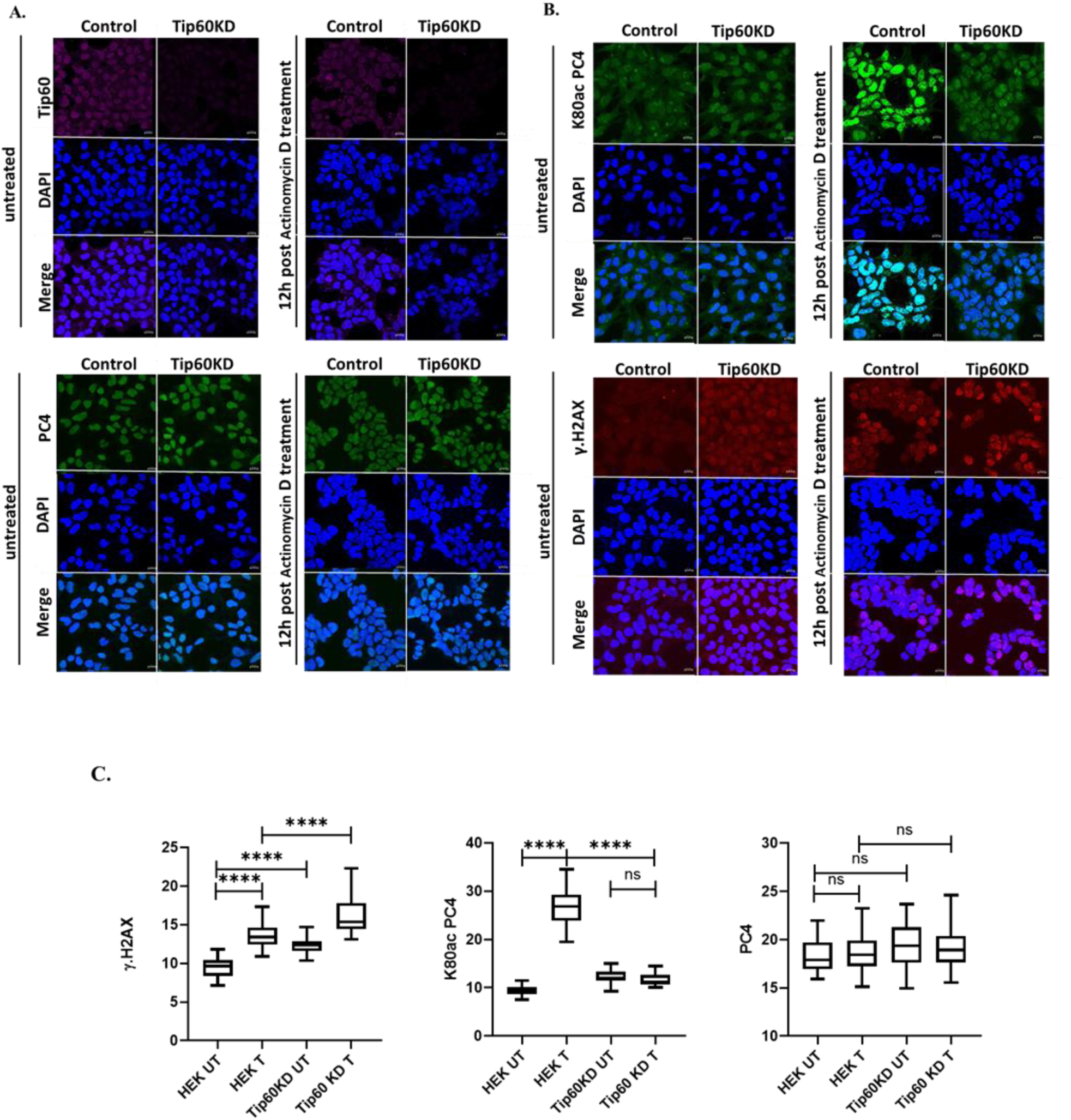
Tip60 is essential for PC4 mediated DNA repair mechanism. (**A, B**) HEK293 WT and Tip60 KD cells were treated 0.2µM Actinomycin D for 12hours. Cells were stained after the aforementioned time point for treatment. Staining with K80Ac PC4 antibody was used to detect the PC4 acetylation level, while PC4 antibody was used to detect the unmodified form of PC4. γ.H2AX was used as a DNA damage marker. (C) Fluorescence intensity was quantified using ImageJ and plotted (n=50 cells). Bars represent mean ± SE; Statistical analysis was carried out by one-way ANOVA using Dunnett’s test for multiple comparisons, \*\*\*\**p* < 0.0001, \*\*\**p* < 0.001, \*\**p* < 0.01 and \**p* < 0.01. (Scale Bar-10µm).

The knocking down of Tip60, drastically reduces the acetylation of PC4, which indicates that the acetyltransferase could be critical for the acetylation of PC4. However, in order to ascertain whether the catalytic activity of Tip60 is responsible for the acetylation, cells were treated with Tip60 specific inhibitor, NU9056. The inhibition of Tip60 acetyltransferase activity upon NU9056 treatment was verified by the H4K8ac levels, which is a known target of Tip60. We found that upon treatment with the increasing concentration of NU9056, there is a dose dependent decrease in the acetylation of histone H4K8 (Fig. 4A, panel 3, lanes 3-6), with the concomitant reduction in the level of PC4K80 acetylation (Fig. 4A, panel 1, lanes 3- 6). However, H3K9ac levels remains unchanged upon dose dependent inhibition of Tip60 catalytic activity, which was used as a negative control.

**Figure 4.**
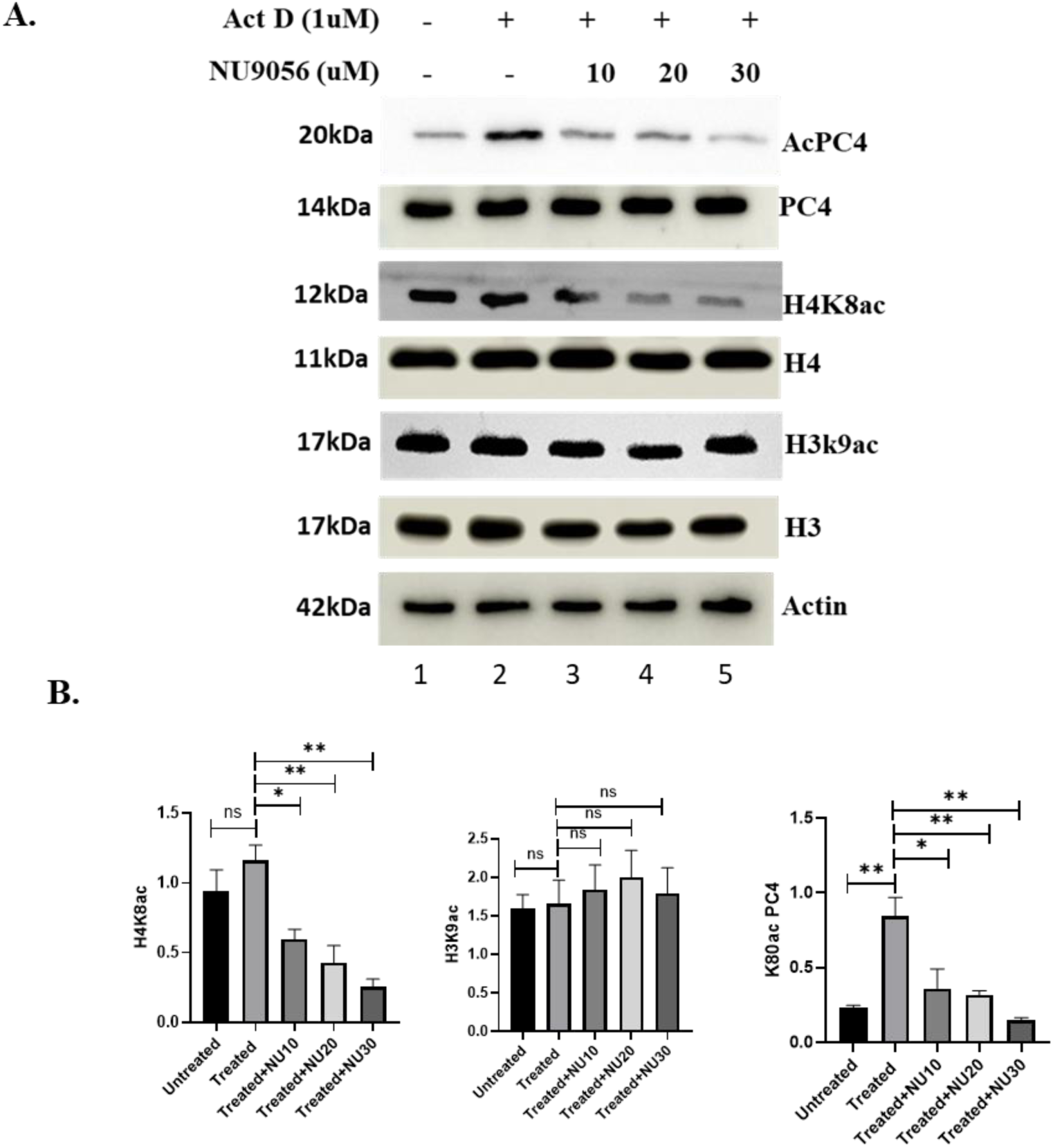
Enzymatic activity of Tip60 is critical for the acetylation of PC4. (A) PC4K80ac expression was checked in cells treated with Tip60 specific inhibitor NU9056 in a dose dependent manner. Decrease in H4K8ac levels is shown as a readout of Tip60 inhibition. The quantification of the dose dependent inhibition of the acetylation has been depicted in the (B). The figure is representative of three biological replicates (n=3). Bars represent mean ± SE; *n* refers to the number to replicates Statistical analysis was carried out by one-way ANOVA using Dunnett’s test for multiple comparisons, \*\*\*\**p* < 0.0001, \*\*\**p* < 0.001, \*\**p* < 0.01 and \**p* < 0.01.

### 3.5. Acetylation of PC4 by Tip60 at K80 site is crucial for its DNA repair ability

PC4 plays critical role in DNA damage repair. In absence of PC4, that is upon knock-down of PC4, γ-H2AX levels gets dramatically enhanced (Fig. 5A, panel 1, lane 2). As expected, transfection of the wildtype PC4 in the knock down cells, significantly, reduced the damage signal, suggesting that indeed PC4 is critical for the DNA repair. Interestingly, the transfection of acetylation mutant PC4, could not reverse the γ-H2AX levels, suggesting that (Fig. 5A, panel 1, lane 3,4) PC4 acetylation at K80 is essential for the DNA repair in the cellular context. In order to visualize the essentiality of PC4K80Ac in cellular DNA repair, we performed the comet assays.

**Figure 5.**
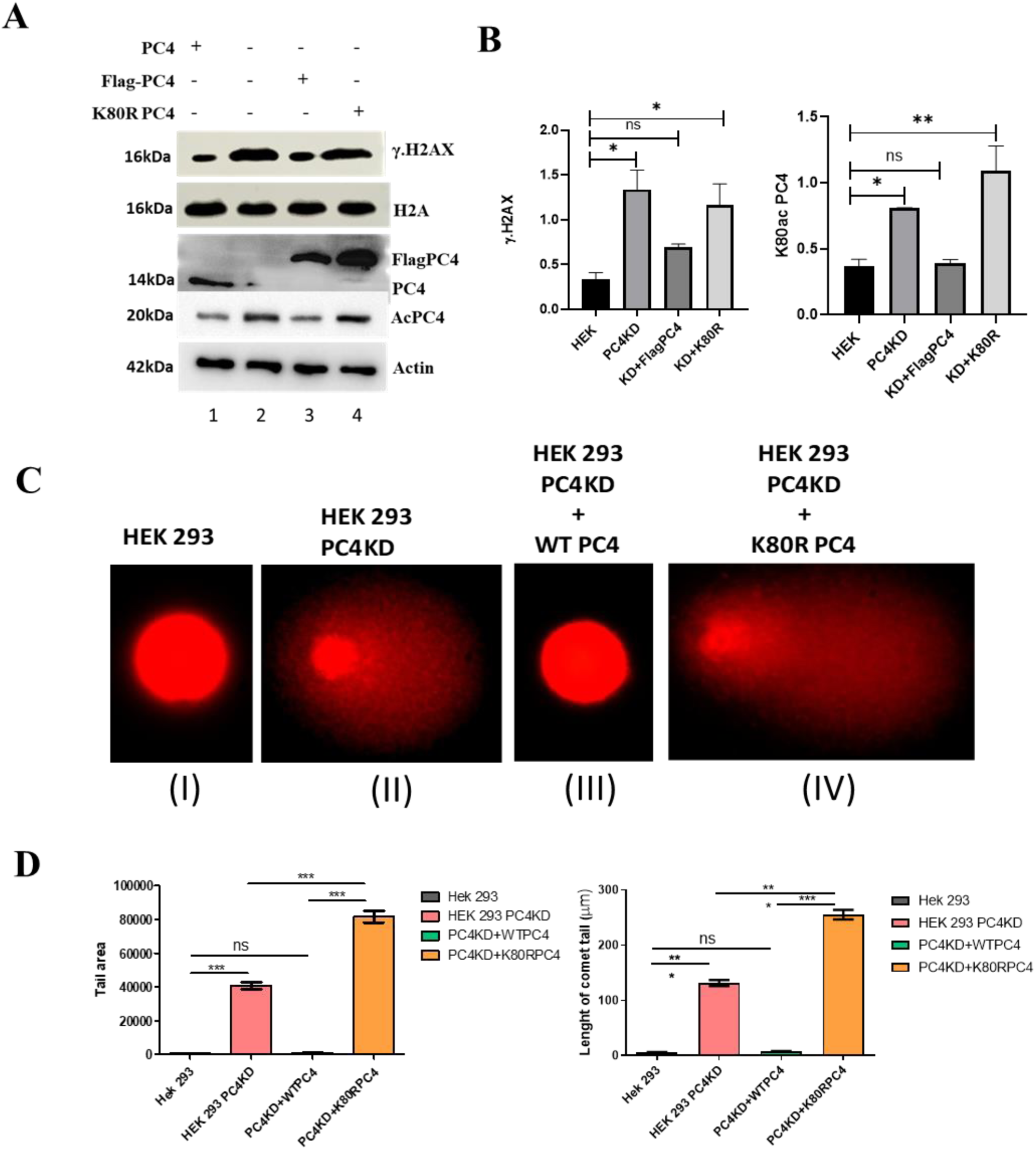
Acetylation of PC4 at K80 residue is critical for DNA repair activity. (A) HEK 293 cells were transfected with indicated plasmids. At 48 hours after transfection, immunoblotting was performed to check the γ-H2AX and PC4 K80Ac expression levels. Representative images are shown. The quantification has been depicted in the (B). The graph shows mean ± SEM; n = 3 for each group C) Alkaline comet assay was performed using WT and PC4 knockdown HEK 293 cells. Representative image of the fields from comet assay obtained from DNA extracted from cells transfected with WT and acetylation defective mutant (K80R) are shown. (D) Comet assay data was quantified by specialized CA software CometScore (150 cells per group). Bars represent mean ± SE; *n* refers to the number to replicates.

In a comet assay, the appearance of the nuclear DNA structures from the DNA embedded in the agarose visualized by fluorescent dye gives an estimate of the extent of DNA damage in the cells. The intensity of the comet tail relative to the head suggests the number of DNA breaks. As depicted in the Fig. 5C, HEK293 cells with PC4, show negligible comet formation, whereas knock-down of PC4, leads to huge comet formation involving more than half of the nuclear DNA (Fig. 5C, panel I versus panel II). Significantly, transfection of wildtype PC4 to the PC4 knockdown cells (Fig. 5C, panel III) completely reversed the comet formation. However, the transfection of the acetylation defective mutant, PC4K80R, could not alter the degree of comet formation (Fig. 5C, panel IV). Collectively, these data suggest that PC4 is critical for DNA damage repair, and mechanistically, it is the acetylation of PC4 by the Tip60, which runs the show.

We have observed that PC4 is downregulated in a substantial number of breast cancer patients. In agreement with this observation, it was noticed that there are a few patient derived highly malignant breast cancer cell lines, where PC4 expression is significantly down regulated [27]. One such cell line is, ZR-75-1, in which PC4 expression is dramatically repressed, due to the over expression of one of the oncomir miR29a. In agreement with our initial data in HEK 293 cells where γ-H2AX levels was enhanced upon PC4 knockdown, in ZR-75-1 cell also, we found γ-H2AX levels to be endogenously high due to low expression of PC4. (Fig. 6A, panel 2, lane 1).

**Figure 6.**
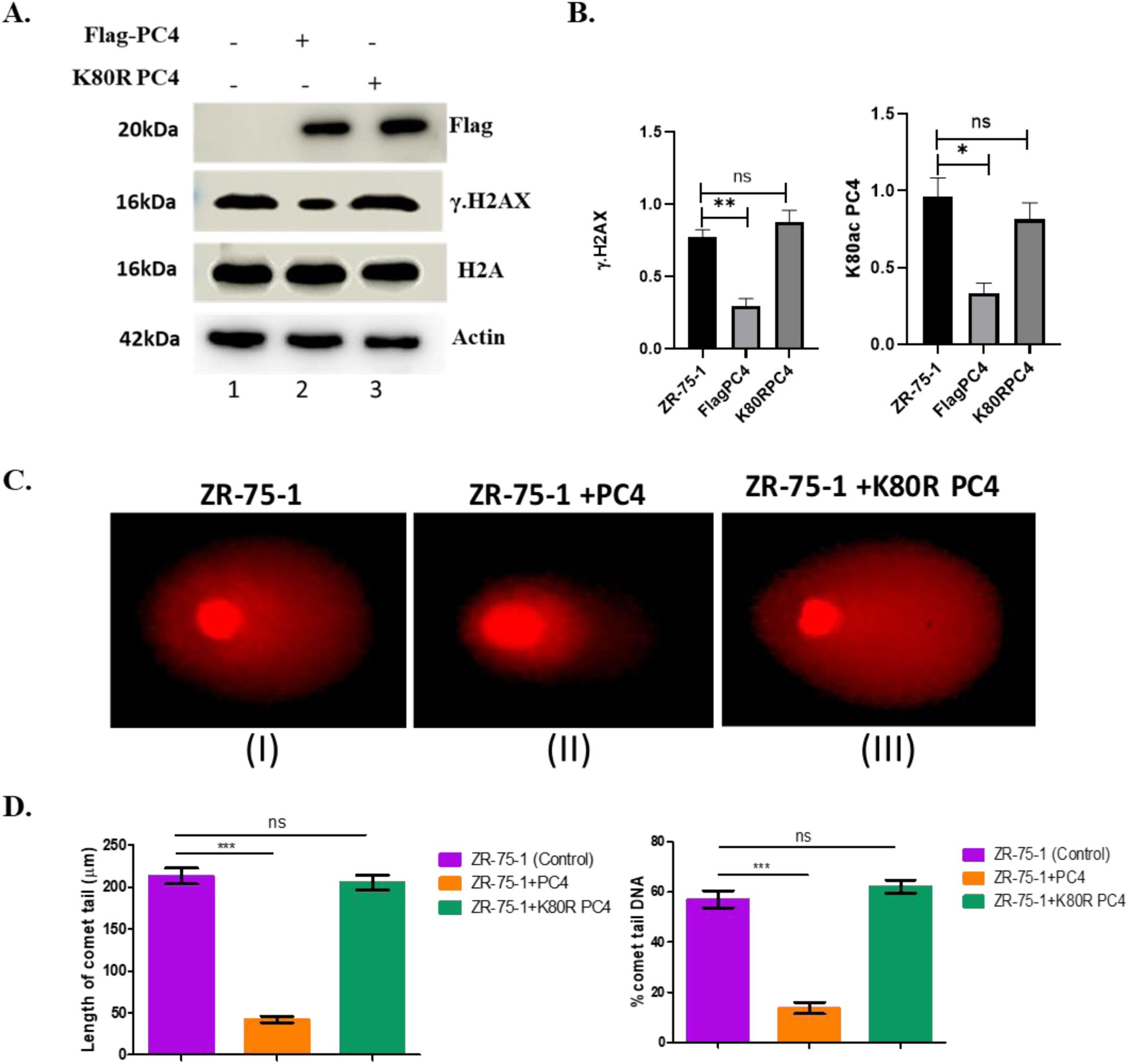
PC4 acetylation is important for DNA repair in ZR-75-1 (A) (A) ZR-75-1 cells were transfected with indicated plasmids. At 48 hours after transfection, immunoblotting was performed to check the γ-H2AX and PC4 K80Ac expression levels. Representative images are shown. The quantification has been depicted in the (B). The graph shows mean ± SEM; n = 3 for each group (C) Alkaline comet assay was performed. Representative image of the fields from comet assay obtained from DNA extracted from cells transfected with WT and acetylation defective mutant (K80R) are shown. (D) Comet assay data was quantified by specialized CA software CometScore (150 cells per group). Bars represent mean ± SE.

**Figure 7.**
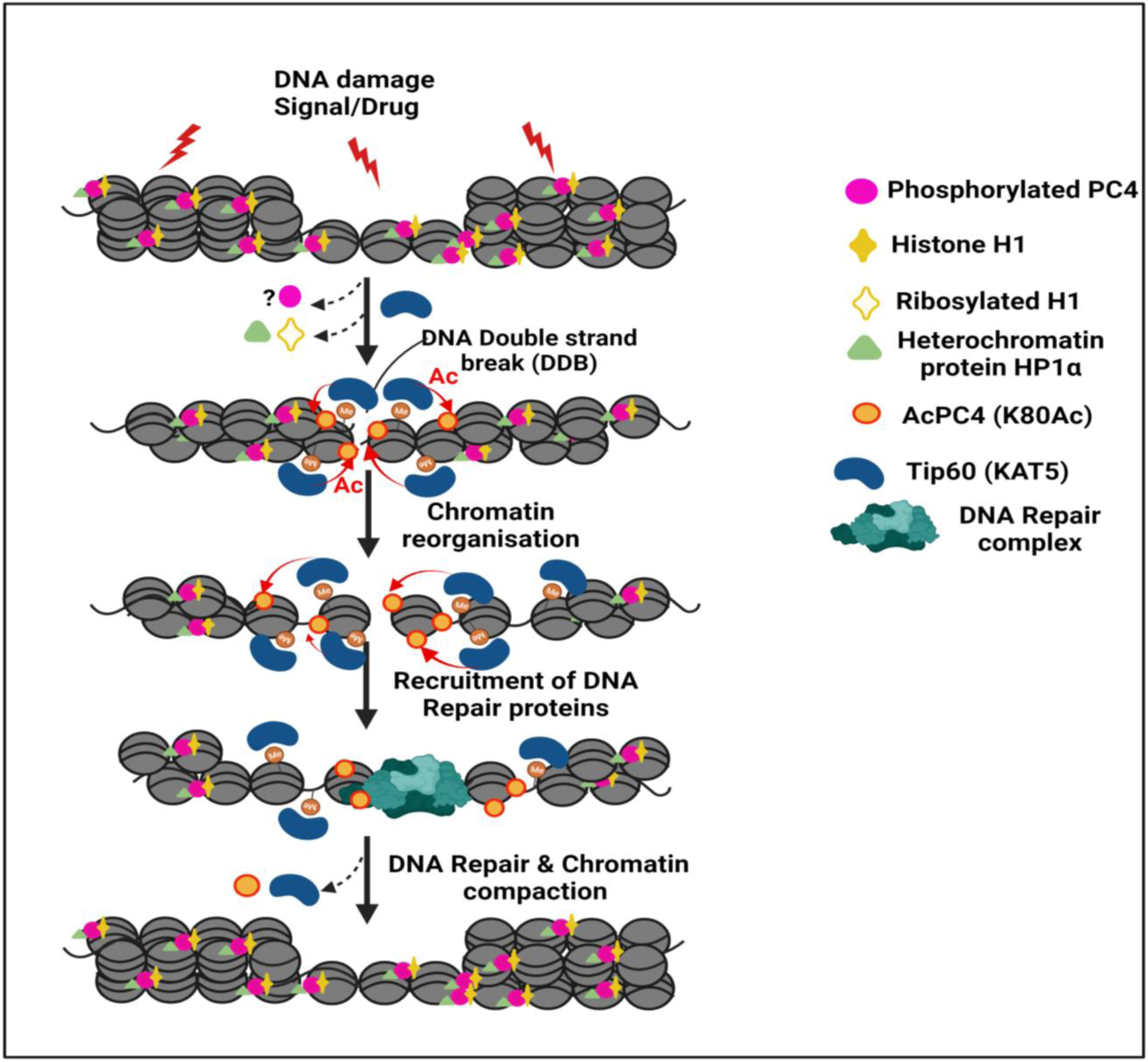
The proposed model of Tip60 mediated PC4 acetylation in response to DNA damage and activation of DNA damage repair. Presumably, upon DNA damage, HP1α gets displaced from damage site along with ribosylated histone H1, allowing Tip60 recruitment and subsequent acetylation of PC4 at K80. These events may lead to chromatin reorganisation, favouring the recruitment of downstream DNA damage response machinery, using acetylated PC4 as one of the interacting partners [2, 28, 29].

Upon transfection with flag-tagged wild type PC4, the phosphorylated H2AX levels decreased significantly in the ZR-75-1 cells (Fig. 6A, panel 2, lane 2) while the acetylation defective mutant failed to reduce the phosphorylation of H2AX. This decrease in the phosphorylation level of H2AX upon PC4 transfection in the ZR-75-1 cells could be attributed to recruitment of functional DNA repair components at the damaged site, by the acetylated PC4 and/or generation of an amenable chromatin environment for the DNA repair. The ability of the acetylated PC4 to assist the DNA repair could be further demonstrated by the comet assay as well. In the ZR-75-1, in which PC4 is endogenously downregulated, shows nucleoids with densely stained tail, which reduces drastically upon expression of wild- type flag tagged PC4 (Fig. 6C, panel II). Fascinatingly, such tailed nucleoid structure reappeared when transfected with a flag construct of PC4 which failed to undergo acetylation FK80R (Fig. 6, panel III). These observations reinforce the fact that PC4 plays a critical role in DNA repair activity for which its acetylation by Tip60 at K80 site is crucial.

To investigate the molecular mechanisms behind the acetylated PC4-medaited DNA damage repair, we performed immunoprecipitation followed by mass spectrometry in HEK293 cells, with and without treatment with Actinomycin D. The result revealed several repair proteins are interacting with PC4 and AcPC4 when subjected to DNA damage specifically. We identified 14 DNA repair related proteins interacting with PC4 in response to DNA damage but not in untreated condition. Out of these 14 proteins, 5 proteins (i.e., PARP1, USP14, RPA1, EPRS1 and PSD5A) were found to be interacting with K80 acetylated form of PC4 also (Fig. S7).

## Discussion

KAT5/Tip60, an acetyltransferase, of the MYST family acetylates tumor suppressor p53, ATM kinase, DNA repair enzyme and androgen receptor transcription factor other than histones. Due to its role in DNA repair and apoptosis, Tip60 is reported to act as a tumor suppressor especially in Breast Cancer. Tip60 acts as a haplo-insufficient tumor suppressor whose expression decreases during breast cancer development and progression [30]. Considering the role of PC4 as a putative tumor suppressor [31, 32] and DNA repair protein [33], here we investigated the possible link of PC4 and Tip60. It has been shown earlier that p300 acetylates PC4 which is abrogated by phosphorylation [34]. We observe that PC4 gets acetylated at a specific site by the acetyltransferase activity of KAT5 both *in vitro* and *in vivo*. KAT5 is a major enzyme that takes part in DNA repair pathway through acetylation of different substrates that mediate the repair of double strand breaks. The role of PC4 in the double strand break DNA repair by both the pathways namely NHEJ and HR is well documented. These contextual connections between Tip60 and PC4 prompted us to investigate the role of this acetylation in DNA damage conditions. We find that indeed in absence of PC4 the phosphorylation status of H2AX, is dramatically upregulated, suggesting the functional role of PC4 in DNA repair. To investigate the role of the acetylated form of PC4 in DNA repair, we transfected the acetylation mutant protein PC4K80R in a PC4 knock down cell. It was found that in PC4 knockdown HEK293 cell line, the extent of DNA damage could be reverted back upon transfecting the wild-type PC4 but not the acetylation mutant PC4K80R construct, as revealed by the comet assay.

We have reported earlier that PC4 is down regulated in a large number (70%) of breast cancer patient samples [27]. In one of the highly aggressive breast cancer cell lines, ZR-75-1, PC4 amount was found to be dramatically low. In agreement with our previous finding, these cells showed, activation of autophagy, radiation resistance and severe reduction in DNA damage repair, as revealed by accumulation of histone H2AX gamma [35].

PC4 acetylation defective mutant was complemented alongside the wild type protein in PC4 knockdown cell line which showed the property of genomic instability (ZR-75-1). Reduction of γ-H2AX levels coupled with comet assay revealed that PC4 might be mediating the DNA repair activity possibly through Tip60 mediated acetylation of PC4 at K80, signifying the early recruitment of PC4 at the DNA damage site [17].

It is to be established whether the acetylation is involved in the recruitment of PC4 and other DNA repair machineries in the damage site and thereby facilitate the DNA repair in mammalian cells. Our initial mass spectrometric data suggest that indeed acetylated PC4 interact with the critical DNA repair proteins, only upon DNA damage (Fig. S7). However, the role of the DNA damage repair favorable reorganization of the chromatin upon Tip60- mediated acetylation of histones and PC4 cannot be ruled out.

## Acknowledgements

This work was supported by Programme Support on ‘Chromatin and Disease’ Grant No. BT/01/CEIB/10/III/01). S.S. is supported by CSIR, India. T.K.K. is a Sir J.C. Bose National Fellow. We acknowledge Amrutha Swaminathan and Surabhi Sudevan for technical support in analysing mass spectrometric data and enzyme purification respectively. Special thanks to late Ms. Varsha Singh for her considerable help and support.

