## supplementary for "KAT5 Acetylates Human Chromatin Protein PC4 to promote DNA Repair"

**# Authors contributed equally to this manuscript**

**Running title:** *Acetylated PC4 in DNA Repair*

**SUPPLEMENTARY FIGURES**

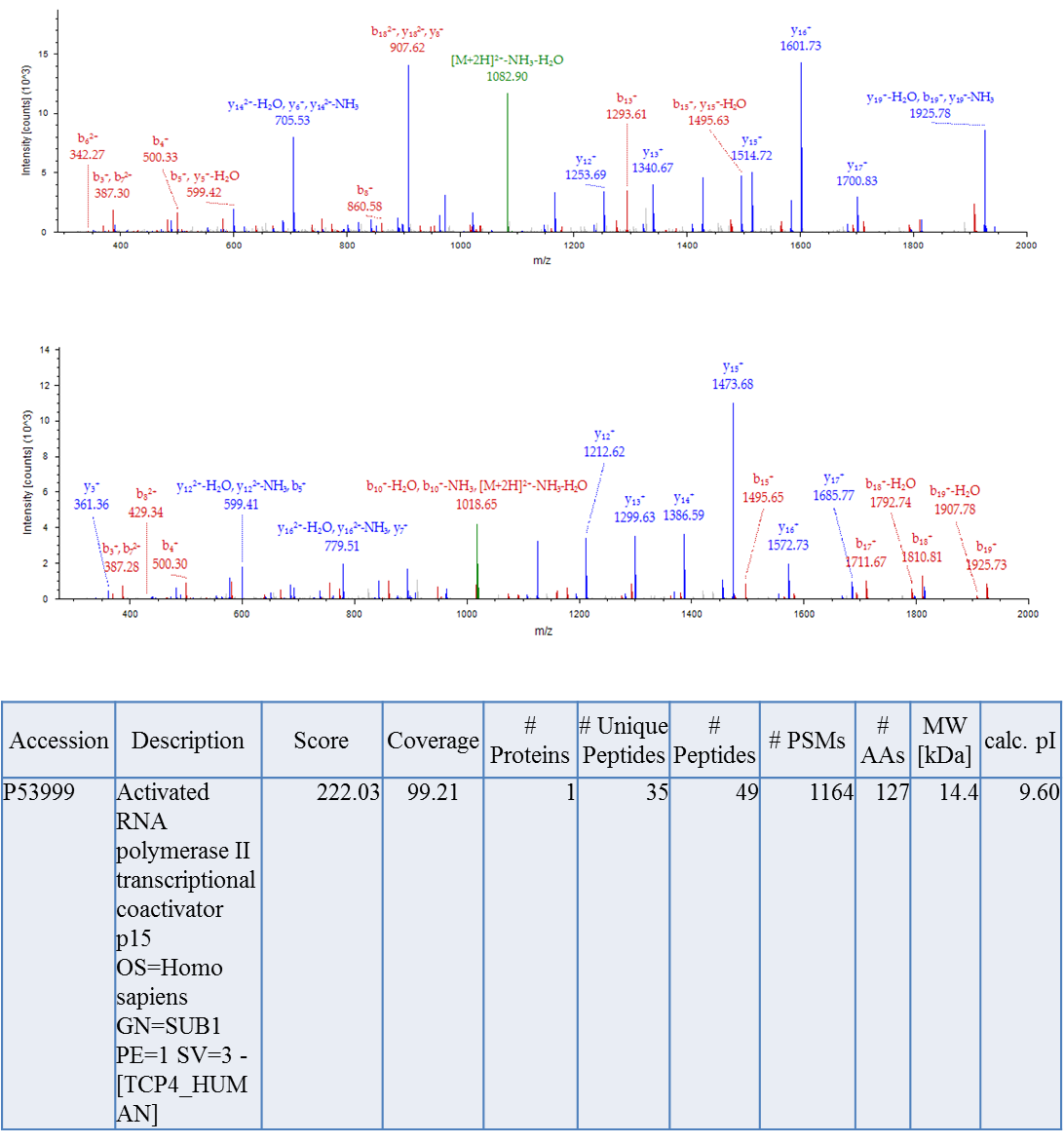

**Figure S1:** Mass spectrometric analysis of *in vitro* acetylated PC4 by KAT5/TIP60: Scoring for the acetylation site identification.

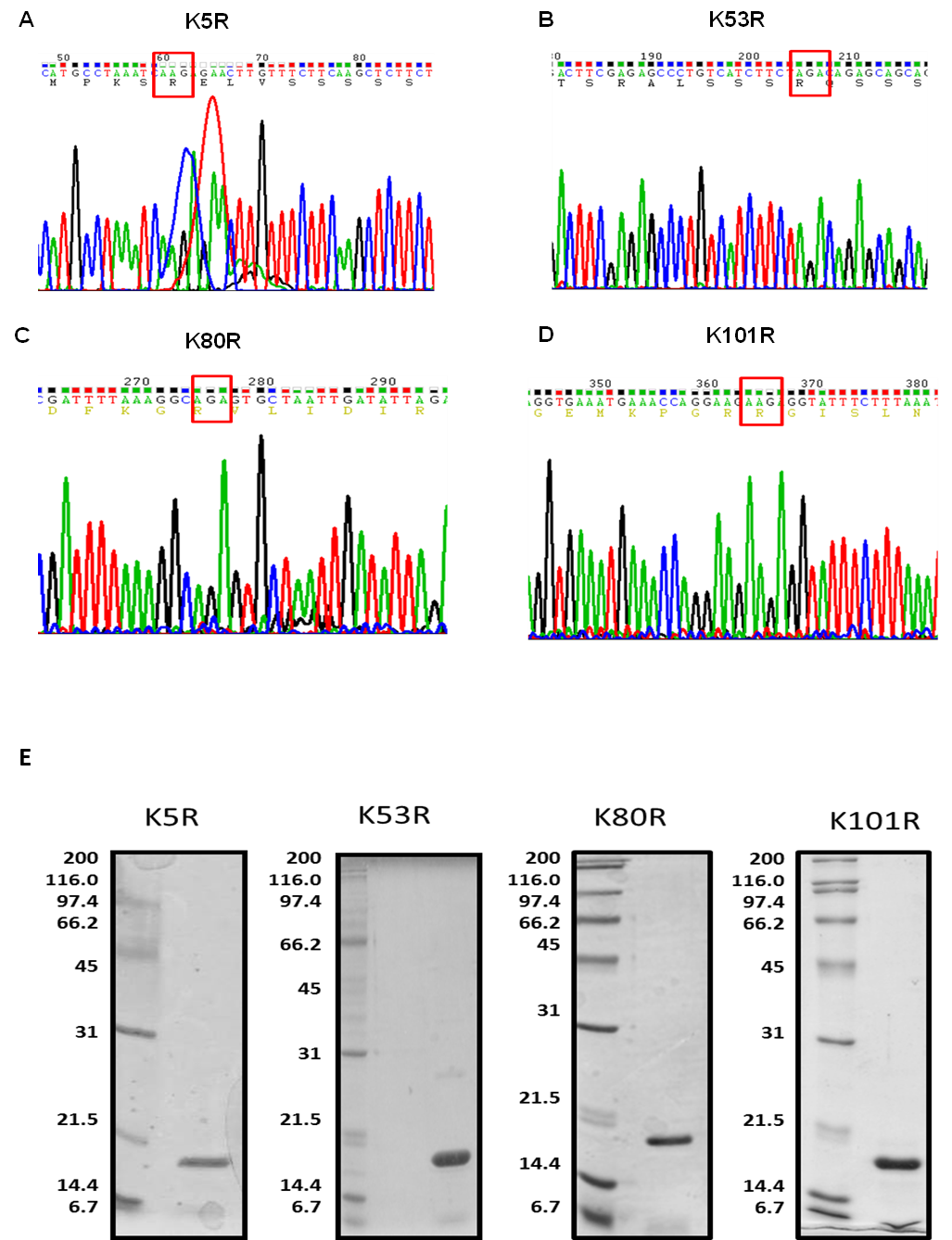

**Figure S2: Purification of bacterial expressed His tagged mutant PC4**: (A-D), Chromatogram representation of all lysine mutants of PC4 generated through site directed mutagenesis. Red square indicates the altered codon to obtain mutant amino acid. (E) Purification profiles of recombinant His tagged PC4 mutants generated by site directed mutagenesis from transformed bacterial culture. His tagged PC4 was purified through a Ni-NTA column. Numbers denote molecular weight in kDa.

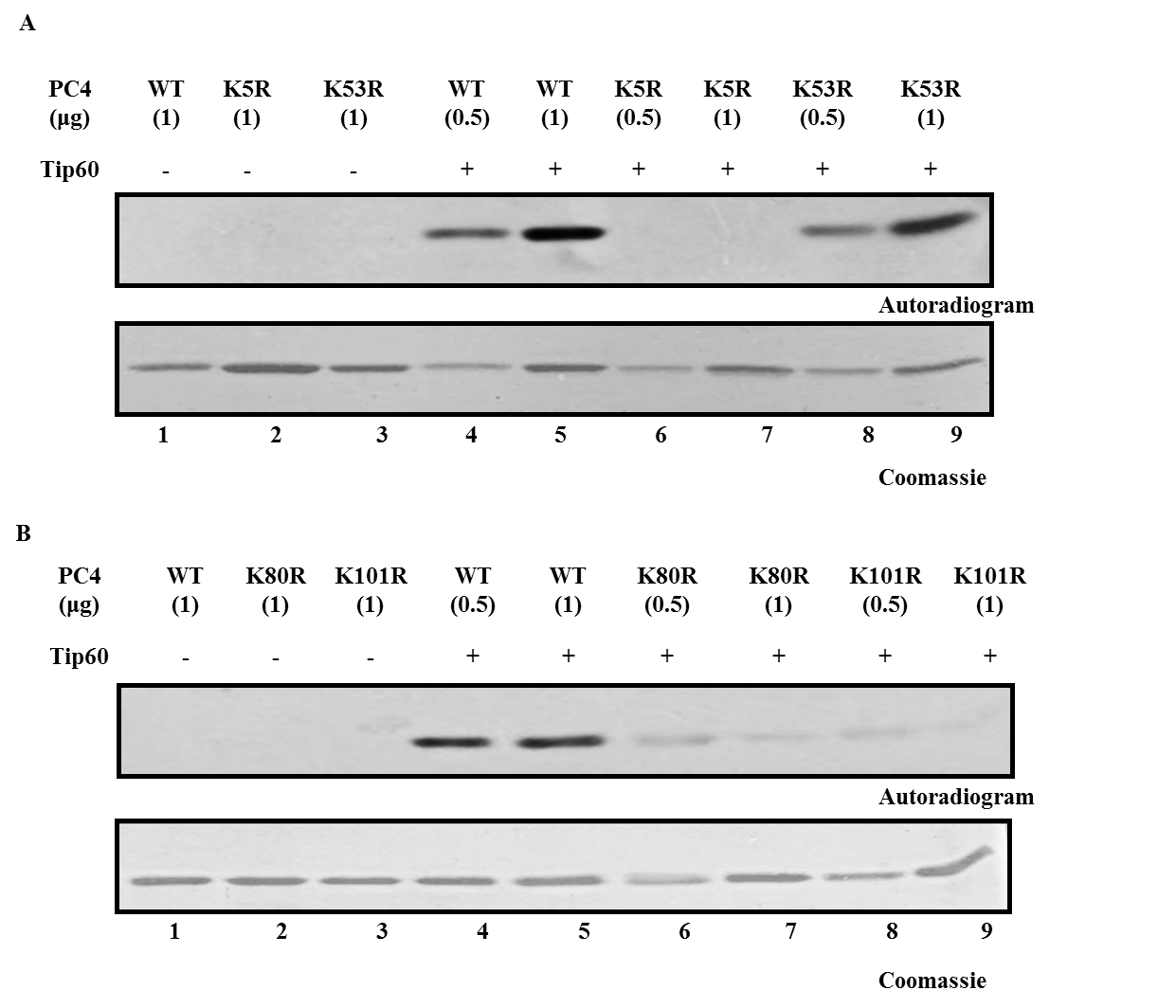

**Figure S3: Tip60 mediated acetylation of lysine mutants of PC4 in a concentration dependent manner**. KAT assay was performed with increasing concentrations of PC4 wild type (WT)

and with increasing concentrations of acetylation site mutants K5R and K53R (Panel A), K80Rand K101R (Panel B) in presence (Panel A and B, lanes 4-9) or absence of Tip60 (Panel A and B, lanes 1-3)

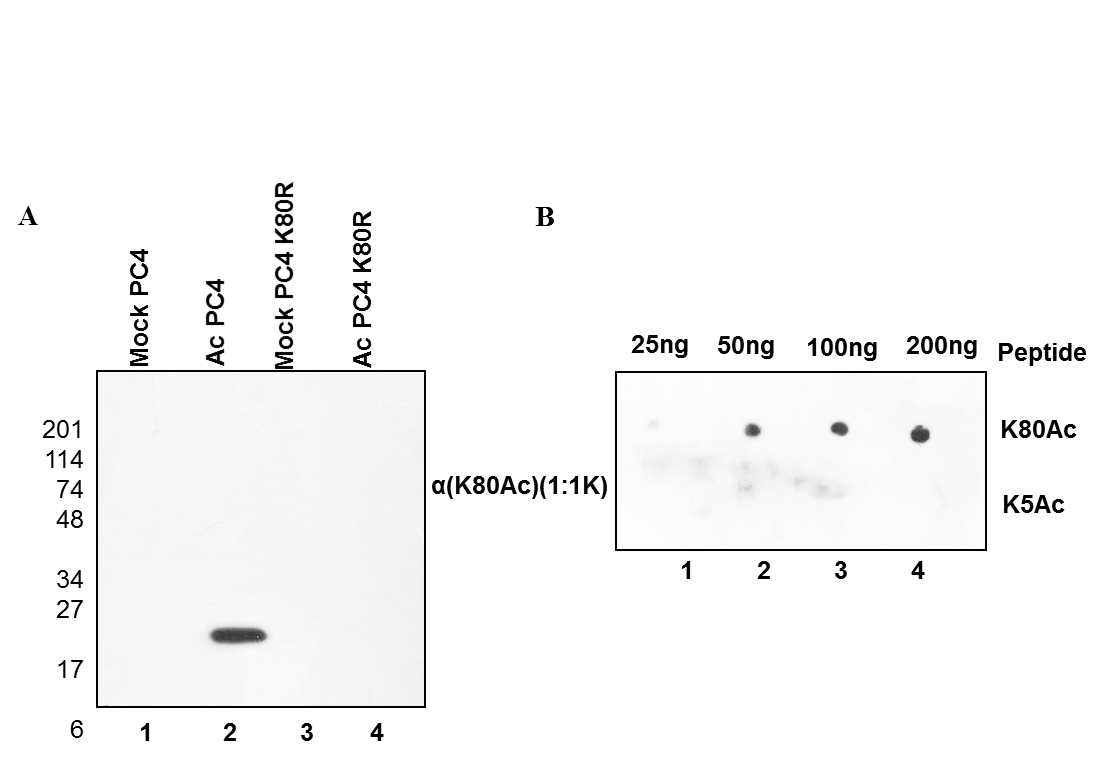

**Figure S4:** (A) Western to determine the specificity of the antibody to recognize the Tip60 acetylated recombinant PC4 (lane 2) over the mock- acetylated PC4 (lane 3). The antibody is specific for K80 site as it failed to recognize the Tip60 acetylated PC4 which had a mutation at K80(K80R) (lane 4). (B) Dot blot to determine the specificity and sensitivity of the polyclonal anti-K80ac antibody (upper panel). To determine its cross-reactivity it was used with another acetylated peptide spanning the K5 site of PC4.

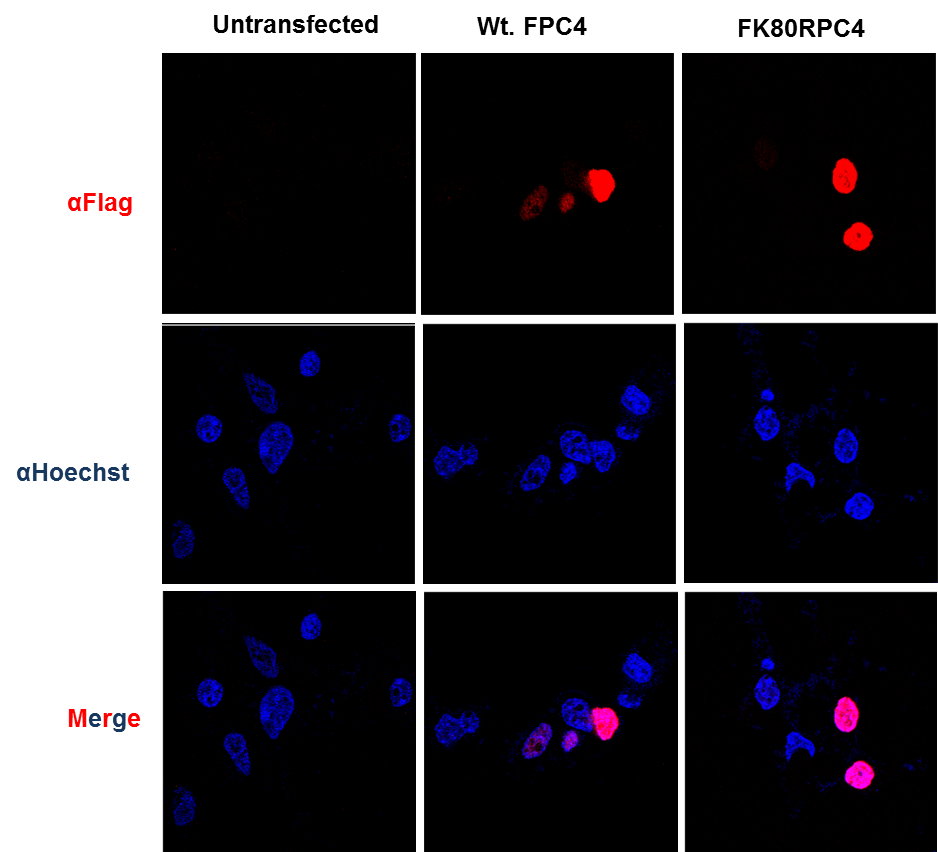

**Figure S5: Cellular localization of acetylation defective mutant of PC4.** Flag constructs of Wild type PC4 as well as acetylation defective mutant PC4 K80R was transfected in PC4sh8 KD cells and immunofluorescence was done with anti-flag antibody to check for its cellular localization. Hoechst was used for nuclear staining.

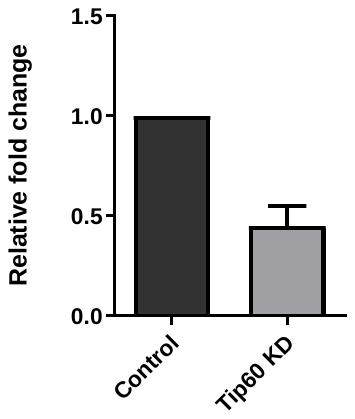

### Figure S6: Quantification of RT-PCR analysis of relative Tip60 mRNA levels in HEK 293 cells transfected with scrambled (control) or Tip60-specific shRNA (Tip60 KD)

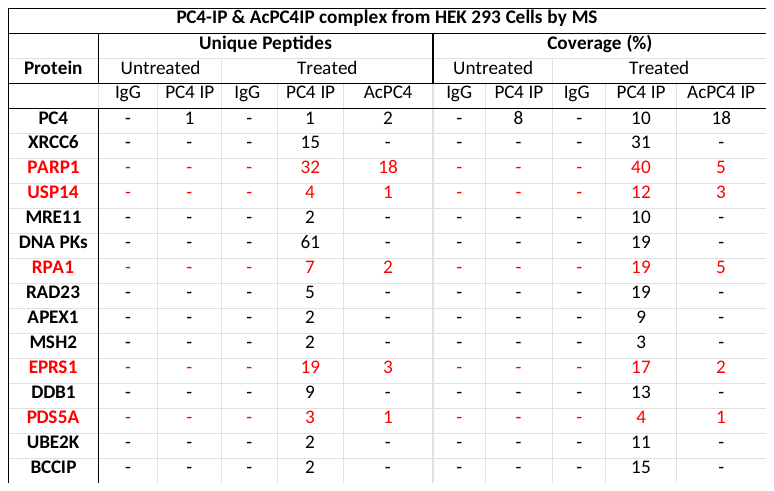

**Figure S7:** Mass spectrometry analysis of the differential binding proteins of PC4 and AcPC4 in HEK 293 cells treated with Actinomycin D. The parameters of the DNA repair protein candidates are shown. The proteins indicated in red are the once interacting both with PC4 and AcPC4 upon DNA damage.
